# Modeling of Rate Heterogeneity in Datasets Compiled for Use With Parsimony

**DOI:** 10.1101/2024.06.26.600858

**Authors:** April M. Wright, Brenen M. Wynd

**Affiliations:** Department of Biological Sciences, Southeastern Louisiana University, Hammond, Louisiana, 70403, USA

**Keywords:** Morphology, fossils, Bayesian, Among-Character Rate Variation

## Abstract

In recent years, there has been an increased interest in modeling morphological traits using Bayesian methods. Much of the work associated with modeling these characters has focused on the substitution or evolutionary model employed in the analysis. However, there are many other assumptions that researchers make in the modeling process that are consequential to estimated phylogenetic trees. One of these is how among-character rate variation (ACRV) is parameterized. In molecular data, a discretized gamma distribution is often used to allow different characters to have different rates of evolution. Morphological data are collected in ways that fundamentally differ from molecular data. In this paper, we appraise the use of standard parameters for ACRV and provide recommendations to researchers who work with morphological data in a Bayesian framework.

## Introduction

It has long been observed that different characters in phylogenetic datasets evolve at different rates (Fitch and Margoliash, 1967a,b; Mayr, 1965; Camin and Sokal, 1965; Kluge and Farris, 1969). A given site or character is subject to a variety of evolutionary forces that may result in either stasis or change. Purifying selection or evolutionary constraint may lead to a character being invariant or changing very slowly. On the other hand, characters of ecological relevance may change frequently in a group of interest.

Often called among-character rate variation (ACRV, or among-site rate variation in the case of molecular phylogenetics), modeling this phenomenon has been of interest since the early days of phylogenetics (Uzzell and Corbin, 1971; Kocher and Wilson, 1991; Irwin et al., 1991). In the early days of molecular phylogenetics, it was appreciated that if all characters had the same rate of evolution, then the distribution of the numbers of changes in a set of characters should be roughly Poisson-distributed. In testing this idea, Fitch and Margoliash quantified changes across a variety of organisms for the gene Cytochrome C, observing that this property only held if invariant characters and hypervariable characters were removed from the dataset (Fitch and Margoliash, 1967b). Various authors tried to incorporate this information in phylogenetic analyses in a variety of ways. Early attempts including parameterizing the character evolution model such that there are invariant and variant sites (Hasegawa et al., 1985; Hasegawa and Kishino, 1989), and using a continuous distribution of evolutionary rates among sites (in a thesis chapter Yang (1992), but subsequently publicized via Yang (1993, 1994)). In 1994, Ziheng Yang proposed the way we predominantly incorporate ACRV in our analyses today: using a discretized gamma distribution. Under this way of modeling ACRV, the rate of evolution for any particular character is drawn from a distribution, often a gamma distribution. For computational tractability, this gamma is treated as discrete, and broken into a number of rate categories. A set of example gamma distributions are shown on Figure 1. The gamma is very flexible, possibly taking on many shapes. Commonly, the gamma distribution will be broken into four categories. These are often considered characters with a slow rate of evolution, characters with a very fast rate, and two intermediate rates. This allows for the evolutionary rates of sites to be more adequately modeled, rather than assuming that they are all share a single rate of evolution.

**Fig. 1.**
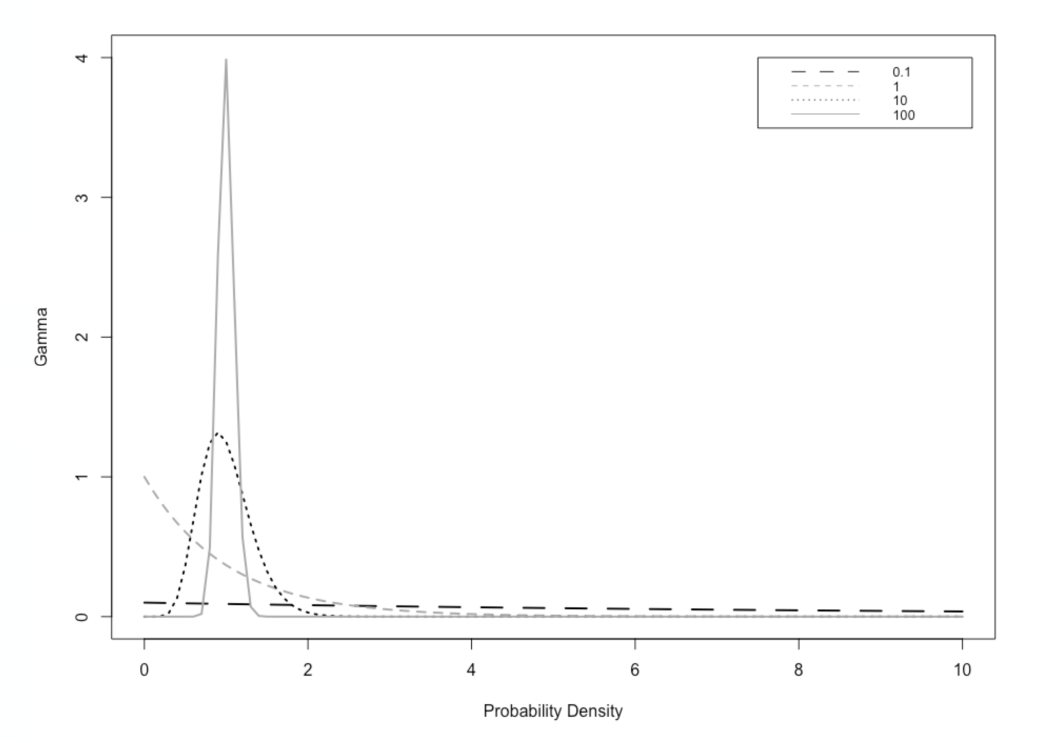
This figure shows several different Gamma distributions. They differ in the value of their *α* value.*α*. While the gamma distribution has two parameters, the shape and the rate, in phylogenetics, the rate parameter is typically normalized to one. This is to aid with interpretation of branch lengths. Thus, *α* is typically the only value estimated in most phylogenetic software. A low *α* will correspond to a very spread out gamma distribution. This means many characters are very different from each other in their rates of evolution (i.e., high ACRV). A high alpha will concentrate characters around the mean rate (1), meaning little ACRV.

Description of a discretized gamma distribution for ACRV was carried out with molecular data. The manner in which molecular data is collected is greatly different from morphological data. The overwhelming amount of research into the understanding of systems like DNA has lead to those systems being viewed as objective forms of data. Choice of marker for a molecular phylogenetic analysis can lead to overall distributions of rates of evolution that differ from other markers, a fact which has been recognized for decades (Cavalli-Sforza and Piazza, 1975). Within any one marker, whether a gene, a restriction-associated site, or another element, there is still rate variation among individual sites and the researcher is likely unaware *a priori* of which sites evolve at which rates, though they may be able to make educated guesses based on codon position and other factors. Where a molecular analysis would see a nucleotide’s position and its identity (ATCG), a morphological analysis would see an array of features that the investigators found most prudent, and the various states in which that feature is expressed across the sample, which introduces bias and subjectivity. Because it can be difficult to capture the complexity of any given feature, morphological phylogenetics often relies on discretizing characters into the various states that are observed in a sample of close relatives (e.g., see Supplement 1 in Ezcurra (2016)).

This contrast between molecular and morphological data becomes evident when exploring paleontological data. For the fossil record, DNA is often inaccessible and so paleontologists have gravitated toward cladistic methods, where the messy nature of paleontological data could be accommodated more flexibly, albeit subjectively. The cladistic model focuses on a unit of measure known as a ‘character,’ which is then discretized into constituent character states that are expressed mathematically as integers. In paleontological studies, the character generally refers to a feature of a structure (e.g., the fourth trochanter of an archosaurian femur), but it is the character state that describes how that feature could be differently expressed among individuals in the study. In our case of the archosaurian femur, there are two general approaches: if the feature is only present in a subset of the sample, the character states will most likely be absent or present, generally represented by 0 and 1 in analyses, respectively (though apomorphic losses would likely be coded as the inverse). If the character is represented in the sample by multiple unique and differentiable forms, then the character is more likely to be expressed as multistate, with additional states (e.g., 2, 3, …, n) representing the various forms of morphological expression in the sample. These characters are modeled in the phylogenetic analysis by transition matrices that describe the expected behavior of the character based on expert assessment, and in cladistics, the null assumes that the rate of change between all states for a character are equivalent. Herein lies the subjectivity that proliferates as a core issue: that no two characters are or could ever be equivalent to one another. With DNA, base pairs follow strict biological rules of organization, which results in quantifiable distinctions between the rates of transfer among or between purines and pyrimidines. Thus, each site (character equivalent) for a given taxon in a molecular analysis shares some degree of comparability that is reinforced by decades of study. This is in stark contrast to the morphological model, where we lack the foundational work that outlines how, for example, different muscle scars appear on different bones at either similar or different rates. Then compound that with the fact that some characters may not even represent muscle scars, but could be tooth morphology or even the presence or absence of an entire bone. This inherent subjectivity in what appears to change to the researcher, or which features capture the morphological underpinnings that support the monophyly of a group, is that in a matrix of morphological characters, they do not each represent the same base unit of measurement. Much as physicists simplify the true complexity in nature by representing the speed of light as the speed of light in a vacuum, or even more simply as just ‘c,’ to ease their calculations (among many other examples), morphologists have stripped the away objectivity and nuance of the skeleton into zeroes, ones, and steps.

The cladistic model relies on the character, and its various states, and most frequently presents results in the form of ’steps’, which is the cumulative total of character transitions required to produce a given topology and where the lowest obtained value is fundamental to selecting sampled trees. This focus on steps and finding the most parsimonious solution (Occam’s razor), often leads to the identification of features that are both variable and thought to be representative of a group. But this potentially introduces an issue, as it is often assumed that traits that evolve quickly may be more evolutionarily labile, and thus should not be as impactful in the model outcome as a trait that is predicted to be more stable because it evolves relatively few times (Cummins and McInerney, 2011; Brocklehurst and Benevento, 2020).

We would not assume that a collection of disparate traits from all across the body would all evolve at the same rate as one another, but this is the default assumption when collecting data assumed to be parsimony informative. Importantly, this is not to the fault of the character nor the researcher, but the prevailing model failing to account for the variation that is ubiquitous in nature. Furthermore, if the inherent structure of the analysis assumes equivalent rates of change between character states, perhaps that also influences the researcher to abandon characteristics that are either seen as too variable to be informative or to write character states such that the more subtle variation is lost, as a means to reduce the size of a matrix. By allowing for there to be variable rates among characters, it is possible to return some of the objectivity to these data, and incorporate our understandings of evolution into the parameterization of our models.

Given these differences, one might ask what ACRV means in the context of a morphological matrix. If workers exclude *a priori* characters that do not vary, or vary in a non-informative way (i.e. apomorphies), what is the expected amount of rate variation in a dataset? In light of the different data collection methods between fields, a critical reappraisal of ACRV handling may be needed. Phylogenetic methods are always being updated and it is important to reflect on decisions that have become staples in a field. For example, most studies that use ACRV use four categories of the gamma distribution. Why? Predominantly because four categories produced a good likelihood for the data, and further addition of categories did little to improve this (Yang, 1994). In this paper, we intend to buck the trend of historicity and examine the use necessity of ACRV in paleontological datasets. We examine if a set of empirical matrices shows support for ACRV and ask how common ACRV is among morphological datasets. If some datasets do not support ACRV, is it harmful to include ACRV (i.e., to overparameterize) these analyses, and we examine the converse question - if ACRV is favored, but not included, what is the effect? Using a large repository of morphological matrices (Wright et al., 2016), we demonstrate that most morphological datasets do indeed favor ACRV. We also show that among datasets that do not favor ACRV, but ACRV is included in the model (i.e., the model is overparameterized) estimates of the different rate categories simply converges to the categories being mathematically indistinguishable. However, in datasets where ACRV is favored but ACRV is not included in the model (i.e., the model is underparameterized), there are effects on both topology and estimates branch lengths Ultimately, we propose a new standard protocol when exploring morphological phylogenetics, integrating reversible jump Markov Chain Monte Carlo alongside ACRV to help researchers appropriately parameterize their analyses while avoiding harming the analysis from either under- or overparameterization.

## Methods

### Data

We re-used morphological datasets from Wright et al (2016). Following that paper, we used 574 total data sets, ranging from 5 to 279 taxa and 11 to 364 characters (Wright et al., 2016). These datasets come from a variety of taxonomic groups and scales.

### Models

#### Common Analytic Elements

All experiments were held constant outside manipulations of the gamma distribution. We estimated phylogenetic trees using the Mk model (Lewis, 2001), which has been shown to perform adequately under a variety of simulation conditions (Wright and Hillis, 2014). We split up the matrices according to the number of character states. This is to ensure that the size of the Q-matrix is appropriate. We used an exponential prior on branch lengths, which specifies that some may be fairly long, but most are expected to be relatively short. This assumption has been shown to be effective across many use cases (Brown et al., 2010).

Finally, estimations were run for 100,000 generations. Convergence was checked by eye in Tracer for ESS values over 200 and well-mixed MCMC traces.

#### Reversible Jump Markov Chain Monte Carlo estimation

We performed several different estimations in RevBayes (Höhna et al., 2014; Höhna et al., 2016). The first was a reversible jump Markov Chain Monte Carlo estimation. This allows the number of parameters to change in an analysis and presents values for parameters in the analysis that are averaged between models. In this case, the model is a random variable within the analysis (Burnham and Anderson, 2002). In this case, we set up a reversible jump on the alpha parameter of the gamma distribution. An alpha parameter of 1e8 corresponds to no rate variation, following a tutorial by Hohna and May (2024). In this case, all the characters in the matrix are expected to be evolving at the same rate. Any alpha value lower than this shows at least some among-character rate variation. The alpha value is fit via MCMC. Thus, each data set may have its own alpha value that is unique to it. This analysis allowed us to see both how many datasets support ACRV, and what the preferred alpha values were per dataset.

For datasets where ACRV was preferred, we also ran a set of estimations in which it was not included in the analyses. This was to assess the impact of underparameterizing the analysis.

Finally, we also ran a set of estimations in which rjMCMC was used to assess the number of gamma rate categories. We allowed for there to be between one (no rate heterogeneity) and 15 categories under the gamma distribution. This was done to assess whether the rjMCMC can be a viable way of detecting not just the presence of rate heterogeneity and the shape of the gamma distribution, but the number of unique rate classes.

### Analysis and Processing

We processed the MCMC logs in the R programming language using Tidyverse (Wickham et al., 2019) and Base R (R Core Team, 2018). In all estimations, we discarded 25% burnin before performing any calculations.

## Results

### Support for ACRV among datasets

As can be seen on Figure 2A, most datasets (385) do support some amount of among character rate variation. This figure shows the results from the rjMCMC analysis. In this case, this indicates that most datasets do support ACRV (right hand of the plot), with a smaller number supporting values on the lefthand side. Very few datasets were equivocal (between the peaks). For datasets that do favor ACRV, how much ACRV is variable? As shown on Figure 2B, the data are once again bimodal, but the lower mode (little ACRV; right-hand side of the chart; 213 datasets) has more mass than the left-hand side of the chart (172 datasets). The left-hand side of the chart indicates the dataset shows substantial ACRV. Thus, this indicates that for many datasets that favor ACRV, the amount of ACRV is small, but non-zero. Taken together, this means that while many datasets exhibit a decisive support for ACRV, the actual amount in the dataset is fairly small.

**Fig. 2.**
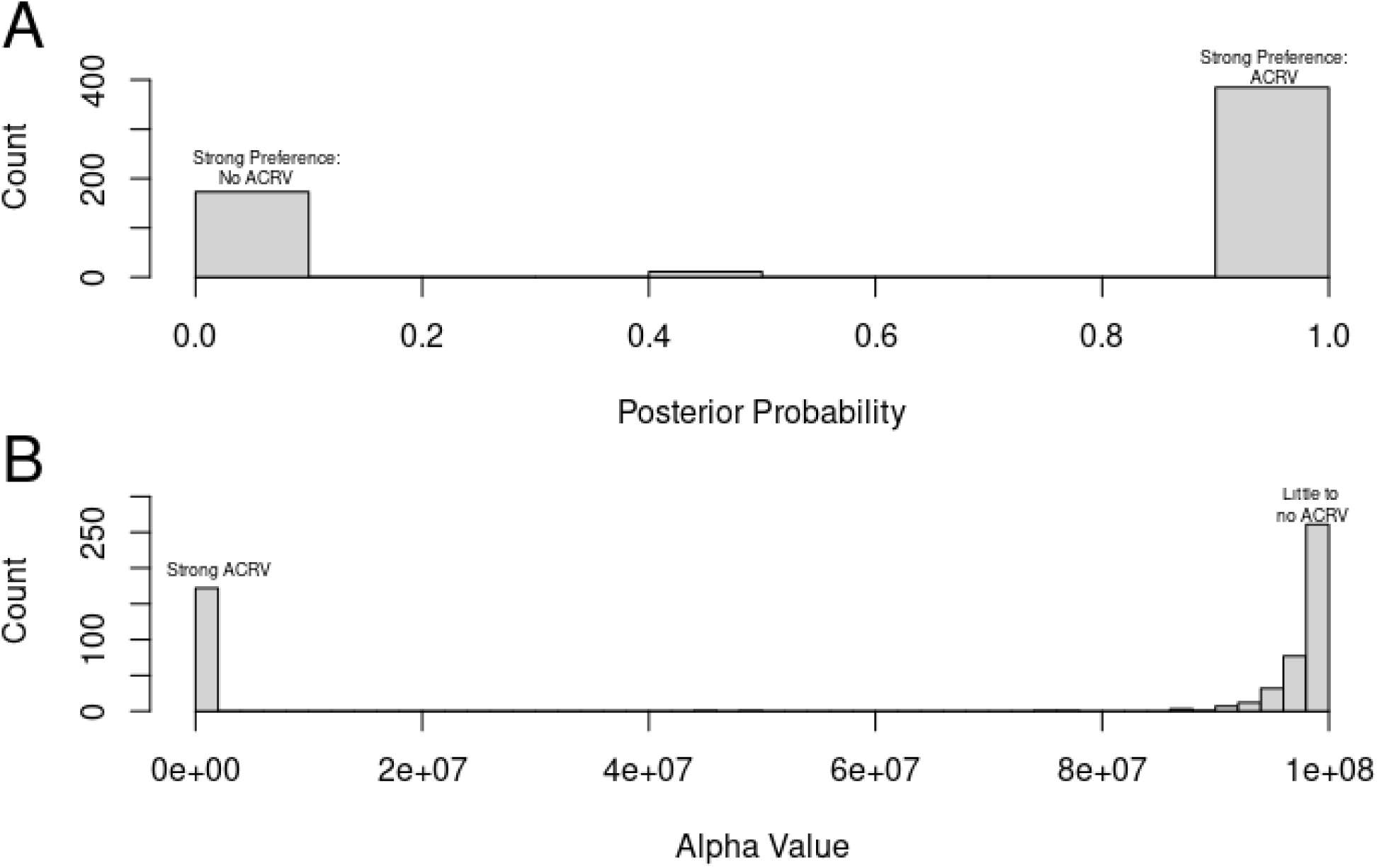
This figure shows the distribution of ACRV preference among our sample.

When we examine the different rate categories, we see similar dynamics. On Figure 3A, we have plotted the four gamma categories. Each color represents a different quantile of a 4-category gamma distribution. Note that one category will always be normalized to one. We can see that for some datasets, there are two distinct low-rate rate classes (red and yellow), the normalized category, and a higher rate class. Though, we also observe that for many datasets, there is substantial overlap between categories (clustering around the normalized rate of one), indicating that, as stated above, while there may be support for ACRV, but the actual amount of ACRV is fairly small. Looking at the net difference between the highest and lowest rate categories (Fig. 3c), for the many datasets that do favor ACRV, we observe that there is a wide range of differences. For a majority of datasets, the difference between the smallest and largest rate categories is close to 0, indicating little ACRV. As indicated on Fig. 2B, there is a minority of datasets for which strong ACRV is supported. The difference between the fastest and slowest rates for these datasets further supports this.

**Fig. 3.**
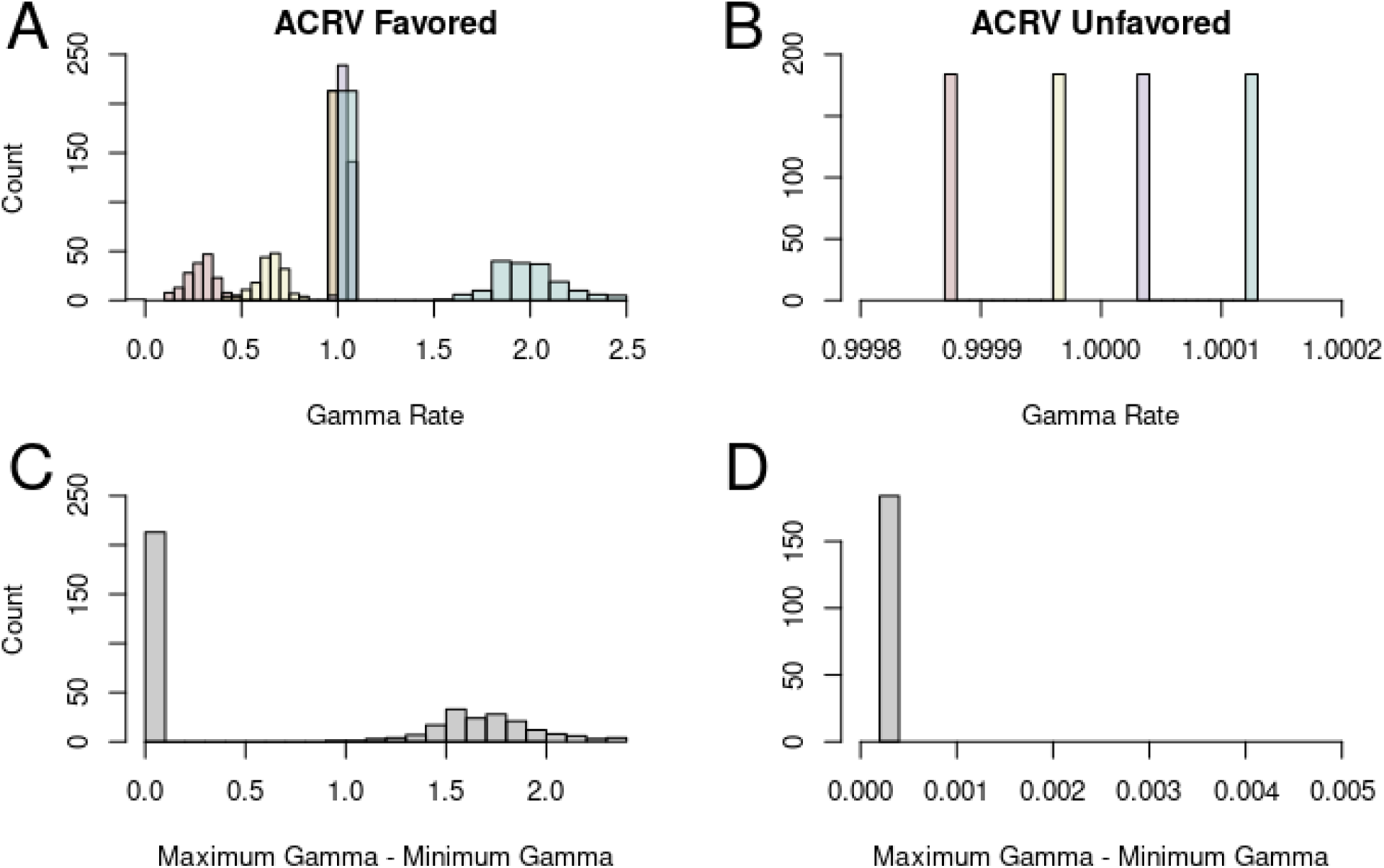
This figure shows the rate variations present in our sample. Site rates are plotted for datasets that A) favor ACRV, and B) do not favor ACRV. A histogram shows the magnitude of difference between the fastest and slowest site rates for datasets that C) favor ACRV, and D) do not favor ACRV

In datasets that do not favor ACRV, the four gamma rate classes all equal approximately 1 (Fig. 3B). The differences between the largest and smallest rate classes for datasets that do not favor ACRV is approximately 0. This indicates that if ACRV is unfavored, the signal of the lack of preference is clearly perceived in these empirical datasets.

### Number of Categories

As can be seen on Fig. 4, datasets that favor the inclusion of ACRV generally favor a larger number of categories than are typically included in phylogenetic analyses. The minimum number of categories favored was 3, with most analyses supporting 7-8 categories, and a smaller number of datasets favored between 9 and 14.

**Fig. 4.**
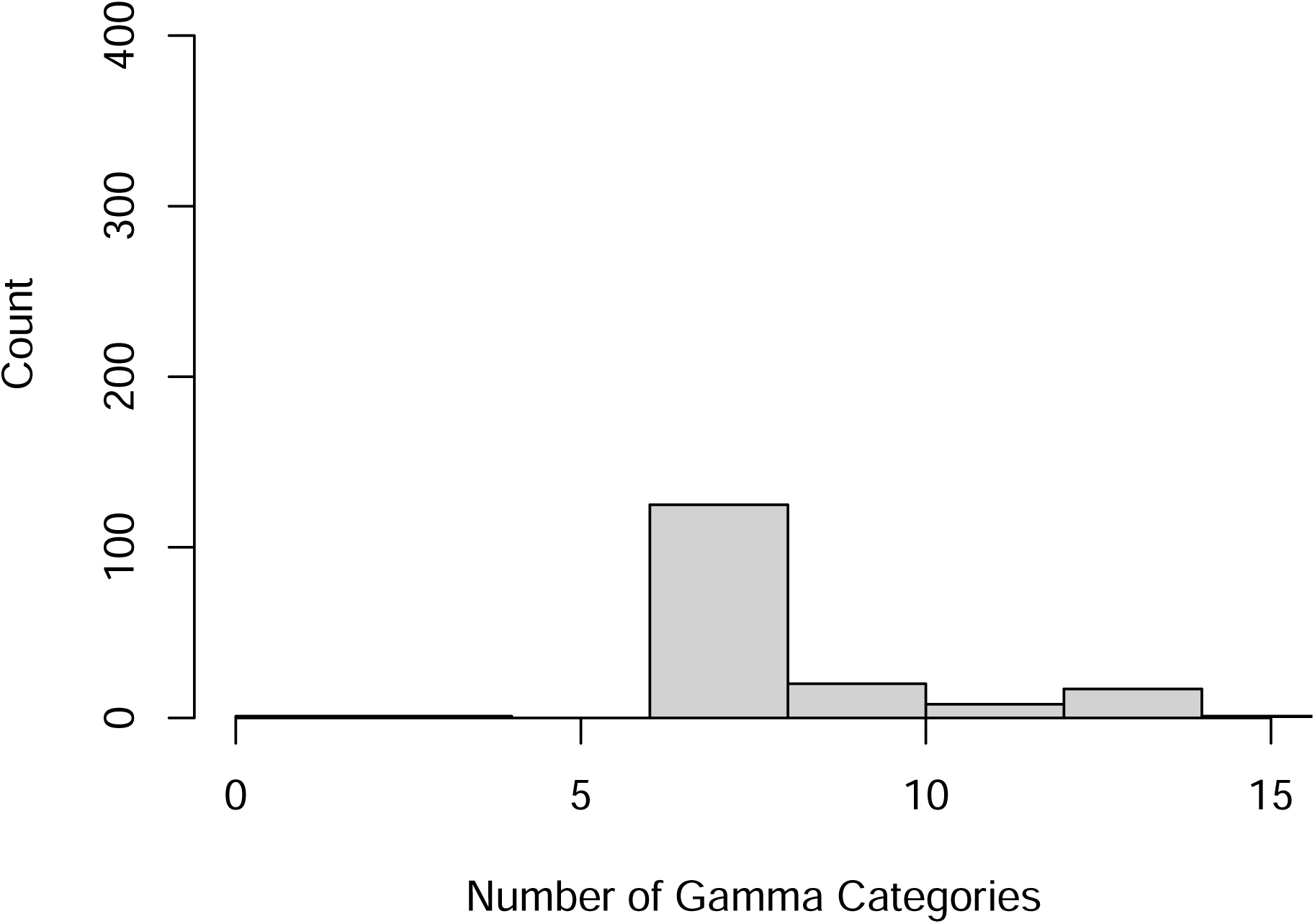
For datasets in which ACRV was favored, we treated the number of ACRV categories as a free parameter and used rjMCMC to estimate its value.

### Consequences of Underparameterizing Models

For datasets in which ACRV was preferred according to rjMCMC, but not included in the model, we see differences between the estimated topologies if it is not included in the estimation model (Fig. 4). For many of these datasets, the Robinson-Foulds distance (Fig 4A) is small, though some estimated trees are nearly entirely different.

To interrogate the branch length differences, we also calculated tree length differences between the rjMCMC trees and those that have no ACRV, and found that trees with ACRV are generally slightly longer. This branch length difference (Fig. 4B) is typically fairly small. We largely expect this pattern when superimposition of changes may be possible along a branch.

## Discussion

### Molecular Models Applied to Morphological Characters

Various authors have raised objections to the applications of molecular models to morphological data (Goloboff, 2014; Assis, 2015). Many of these objections center on the actual substitution process: that the Mk model assumes equal forward and backwards transitions, though other models have been proposed to alleviate this. What has received comparatively less attention are other assumptions of phylogenetic modeling that are not formally part of the Mk model. In this paper, we have assessed the role of gamma-distributed among character rate variation in morphological phylogenetic analyses.

As discussed in the introduction, morphologists collect data in fundamentally different ways than molecular biologists. At the time that the most common methods for dealing with ACRV was published (**?**), molecular sequences were largely being collected by Sanger sequencing, in which the researcher would target a molecule or gene based on its molecular properties (Simon et al., 1994; Hillis, 1996). COI, for example, is a common gene for molecular analyses of relatively close relatives, while housekeeping genes are often used for deeper divergences. No matter what molecule was chosen, the expectation was still that some columns in the alignment will evolve at different rates. Some might be invariant, some will be hyper variable. In today’s world of high-throughput data, variation is still expected. Genetic elements, such as Ultra-Conserved Elements (UCEs) have a low-rate core and higher-rate flanking regions. Thus, among-character rate variation remains critical.

For morphological characters, there is still a pervasive assumption of parsimony informativity in characters. That a character should be diagnostic of a group, changing once or very few times on a tree. If this is true, there should be no real need for ACRV, as all the characters will change at about the same rate. However, this is not always the case, as our results show. As larger datasets become common, issues such as ACRV and character inapplicability become an issue.

Parsimony methods, such as implied weighting take aim at the ACRV issue, by down-weighting characters that are not parsimony informative. Implied weighting incorporates some aspect of rate heterogeneity into the parsimony model, and in doing so, we generally find implied weighting trees to more closely resemble trees estimated from a Bayesian approach, than either do to equal weights parsimony (Goloboff, 2014). This can be shown in the reconstructed histories for the archosauriform clades, Doswelliidae and Proterochampsidae, which have historically been reconstructed as sister to one another in parsimony analyses (Dilkes and Sues, 2009). However, this monophyly was challenged when bayesian approaches were implemented to try and untangle relationships just outside the crown of Archosauria, where the Doswelliidae was reconstructed as a clade within the Proterochampsidae (Wynd et al., 2019). These results have since been supported through implied weights parsimony, suggesting that the inclusion of rate heterogeneity may narrow the gaps between these methods (Ezcurra and Sues, 2021). Parameterizing ACRV using a gamma distribution does not require that any characters are downweighted. All characters in a Bayesian analysis can still be used equally. Characters that are not parsimony informative (i.e. do not favor one tree over another) can still be used to assess rates of evolution and branch lengths - which may, in fact, be informative about tree topologies.

In this exploration, we found that about three-quarters of datasets do indeed support at least some ACRV (Fig. 2A). To the right-hand side of Fig. 2A, the overwhleming majority of samples in the posterior trace for a particular dataset supported ACRV. In an rjMCMC analysis, the proportion of the samples that include ACRV is a proxy for the posterior probability of ACRV. Thus, for these three-quarters of the datasets that support some amount of ACRV, this is a very strong preference. For the vast majority of datasets, ACRV was either supported or denied with extremely high posterior probability. What this means is that for these data, using rjMCMC during the estimation process can lead to fairly high certainty about the presence or absence of ACRV, and the appropriateness of adding it to the model.

For datasets that support ACRV, rate classes are sometimes, but not always fairly distinct (Fig 3A). In Fig. 2B, there is a cluster of datasets to the right-hand side of the plot. These datasets, while ACRV was supported in the rjMCMC analysis, have very little actual ACRV. Many datasets exhibit fairly significant ACRV, with the largest rate classes often being multi-fold larger than the smallest (Fig. 3). Additionally, when the number of categories is a free parameter in the analysis, we generally see that more categories than four are supported (Fig. 5).

**Fig. 5.**
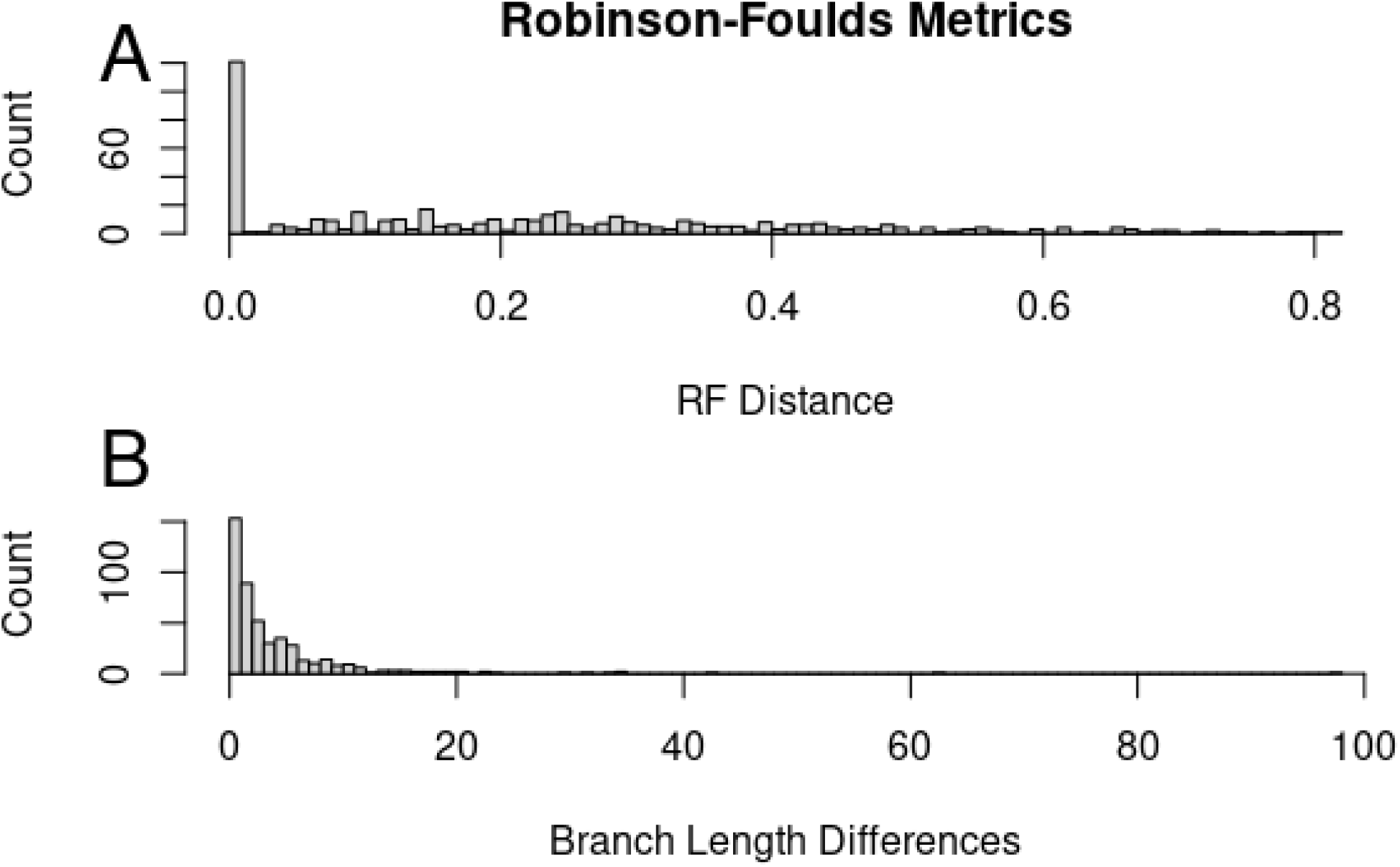
This figure shows the rate variations present in our sample. Site rates are plotted for datasets that A) favor ACRV, and B) do not favor ACRV. A histogram shows the magnitude of difference between the fastest and slowest site rates for datasets that C) favor ACRV, and D) do not favor ACRV

For datasets that did not support the use of ACRV, the estimation converges to a solution in which all four Gamma rate categories are distributed around 1.0, with a less than one-ten-thousandth difference between each rate category (3C), resulting in virtually no difference between the highest and lowest rate categories (Fig. 3D). Thus, if there is truly no ACRV, using an rjMCMC model is not harmful. The signal of there being no ACRV is easily picked up during estimation.

In terms of the estimated phylogeny, for many datasets, the inclusion of among-character rate variation caused the estimation of a different topology. In about 80 of the 375 datasets that supported the use of ACRV, underparameterizing the model made no difference. For the remaining 295 datasets, at least some relationships were different. Likewise, most branch lengths are fairly similar between analyses, but can be quite large for some datasets. This is expected, given that for many datasets, the actual amount of ACRV is small, thus the underparameterization artifact ought to be small.

Our concrete recommendation is that researchers using morphological datasets use a reversible jump MCMC approach. These flexible approaches enable researchers to assess if we need to parameterize ACRV, while simultaneously estimating the parameters of the ACRV model. Parameters reported back are averaged across models, thus taking into account the uncertainty in the model itself. As shown on Figure 2A and 3C, the model easily detects if ACRV is absent and all rate categories are the same. This approach fundamentally allows the researcher to blend the steps of model selection and phylogenetic estimation. Compared to stepping stone model selection, the estimation time is considerably shorter and less resource-intensive. In a benchmark test, rjMCMC took 10 hours and 3 minutes to converge on a single processor, while stepping-stone model slection (Xie et al., 2011) took 24 hours 22 minutes on a single processor.

### Model Averaging, Model Choice, and Complexity

There is a voluminous literature on model selection in phylogenetics (Posada and Crandall, 1998; Huelsenbeck et al., 2004; Posada, 2008; Xie et al., 2011; Baele et al., 2013). Long considered an essential step in the inferential pipeline, if model selection is needed at all when priors are well-defined is now an open question (Guimarães Fabreti and Höhna, 2023). Researchers often wonder when a model is complex enough, when a model is too complex, if it matters, and if models are adequately capturing the generating process of their data (Brown and Thomson, 2018; Mulvey et al., 2024). But an additional question is how can we incorporate the uncertainty in our models? Bayesian model averaging has been a widely used technique in a variety of sciences to answer analytical questions, including systematics (Huelsenbeck et al., 2004; Posada, 2008; Opgen-Rhein et al., 2005; Beaulieu et al., 2013). However, within systematics, it has not been commonly applied for model selection for phylogenetic inference. Model averaging treats the model as a random variable in the analysis and provides a posterior probability of the model itself (Burnham and Anderson, 2002). Distributions of potential values for a given parameter are averaged across different models.

Using reversible jump MCMC and a set of empirical matrices, we aimed to answer a set of questions: How many datasets in morphological data have ACRV, and how much do they have? What we found is interesting on several fronts. Even though characters are often collected with parsimony in mind, they are often not perfectly parsimony-informative and ACRV still exists for most datasets. We also found that many datasets have a great deal of ACRV - often favoring more than four categories. But we also found that this is distinctly detectable with rjMCMC. As shown on Figure 2, very few datasets are at all equivocal about the presence or absence of ACRV. Likewise, averaging over the alpha value doesn’t seem to hurt the inference. For datasets that do not favor ACRV, model averaged parameter estimates for the categories of rate heterogeneity come out approximately equal, indicating no ACRV is present. Thus, blending the model selection and inferential steps does not appear to be harmful for this application.

Being able to take into account and visualize the uncertainty in our estimations is a core benefit of Bayesian estimation. Bayesian model averaging approaches apply this same logic to the model itself, in effect asking not simply what is the best-supported model, but how does the uncertainty about the model impact our estimated parameters? In 2004, Huelsenbeck et al. posited that under a model-averaged framework, “the inference of phylogeny would not depend on any specific model, but would be integrated over uncertainty in the model parameters.” This is a great promise for these methods: many authors have been critical of the Mk model and its assumptions. Model averaging allows us to walk through submodels, interrogating the aspects of each that work for our particular data and reporting solutions that take into account the uncertainty in the generating mechanism of the data. If researchers want to be Bayesians, why not be Bayesians about the whole analysis?

## Conclusions

As we come to understand how best to work with morphological data in a Bayesian context, we hope that researchers will use all available tools to think deeply about both the collective and individual characters and how they should be treated in any given analysis. The sluggish adoption of Bayesian methods among morphologists has lead to a set of gaps in the literature related to basic model functionality. In this publication, we demonstrate that among-character rate variation (ACRV) is very prevalent in morphological data. We also demonstrate that the signal for whether or not ACRV is present is very clear and detectable using reversible-jump Markov Chain Monte Carlo (rjMCMC). Lastly, we recommend that researchers adopt rjMCMC to efficiently estimate if a dataset exhibits ACRV, and how to parameterize it.

## Supplementary Material

Data available from the Dryad Digital Repository: https://datadryad.org/stash/share/db76qKKwJawRNkX00BKYxKmqfuOuIhUd9x2OSK1J5pA. https://datadryad.org/stash/share/db76qKKwJawRNkX00BKYxKmqfuOuIhUd9x2OSK1J5pA.

## Acknowledgements

AMW and BMW were supported on NSF DEB 2045842 and AMW was supported on NSF CIBR 2113425. AMW additionally received funds from Southeastern Louisiana University’s Dyson Fellowship.

